## Supplemental files for "Computational analysis of long-range allosteric communications in CFTR"

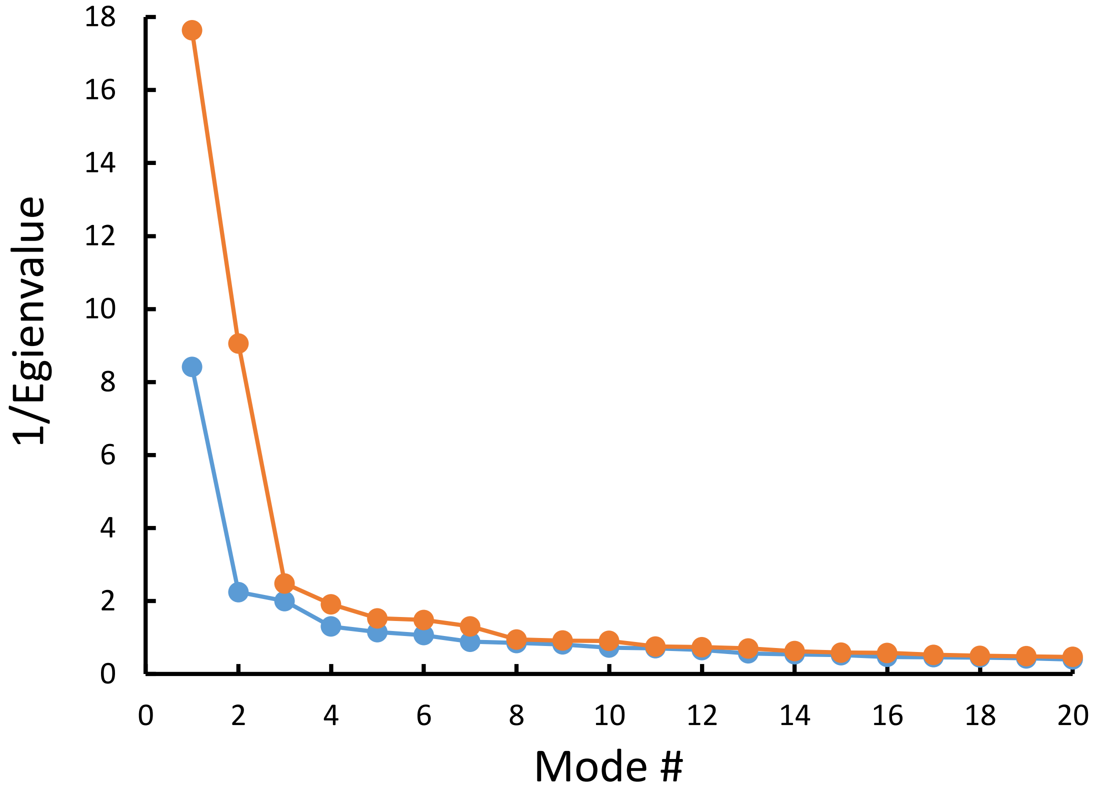


**Figure S1.** Eigenvalue decays as a function of mode number, shown for ATP-free (orange) and ATP-bound (blue) human CFTR (PDB ID 5UAK and 6MSM, respectively).

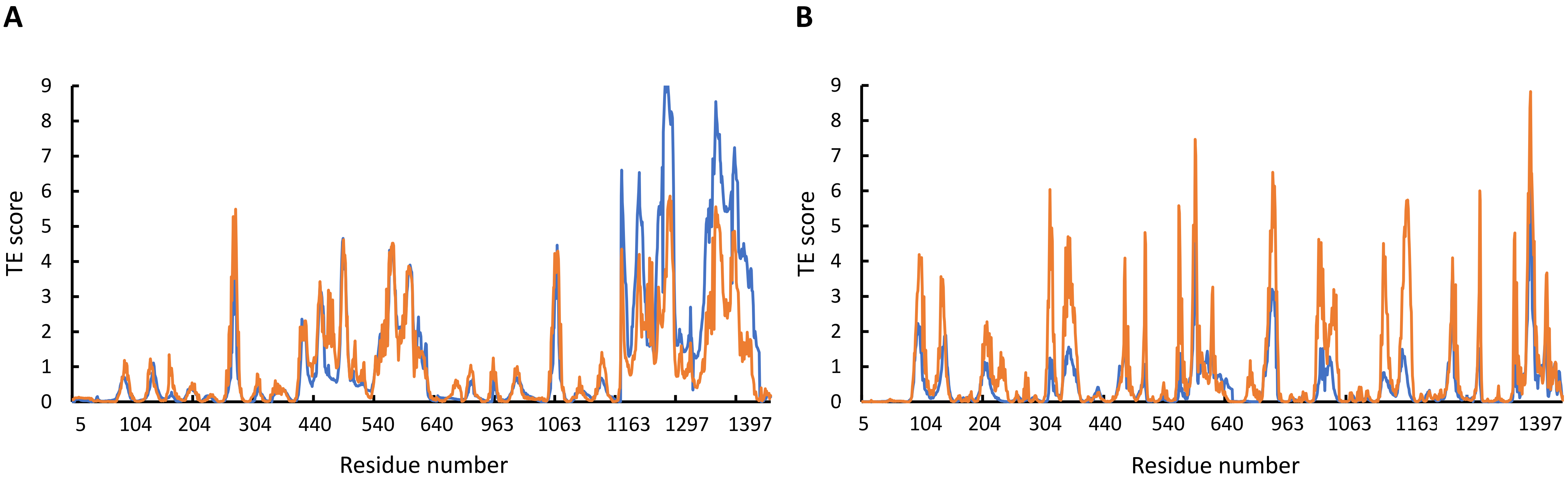


**Figure S2. Sensitivity of the results to Rcut.** Shown is the amount of information transmitted (TE score) by each residue of dephosphorylated ATP-free (A) or phosphorylated ATP-bound (B) human CFTR calculated using Rcut of 7 or 10 Å (blue and orange curves, respectively).

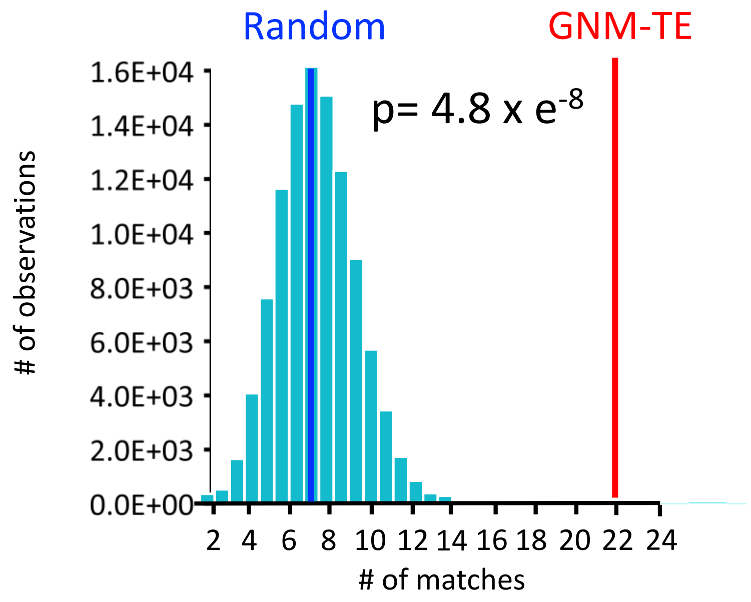


**Figure S3. Co-localization of the GNM-TE peaks with the hinges identified by GNM.** 100,000 sets of 60 randomized positions were generated, and for each such set the number of matches (within a cutoff distance of ≤4 Å) to a position of a hinge residue that was identified by GNM was counted. Shown is the probability distribution for the 100,000 random sets (cyan bars). Blue and red vertical lines indicate the mean value of matches for the random and for the GNM-TE determined peaks, respectively. Also shown is the p value obtained by one-tailed hypothesis tests with a significance level of 0.05.

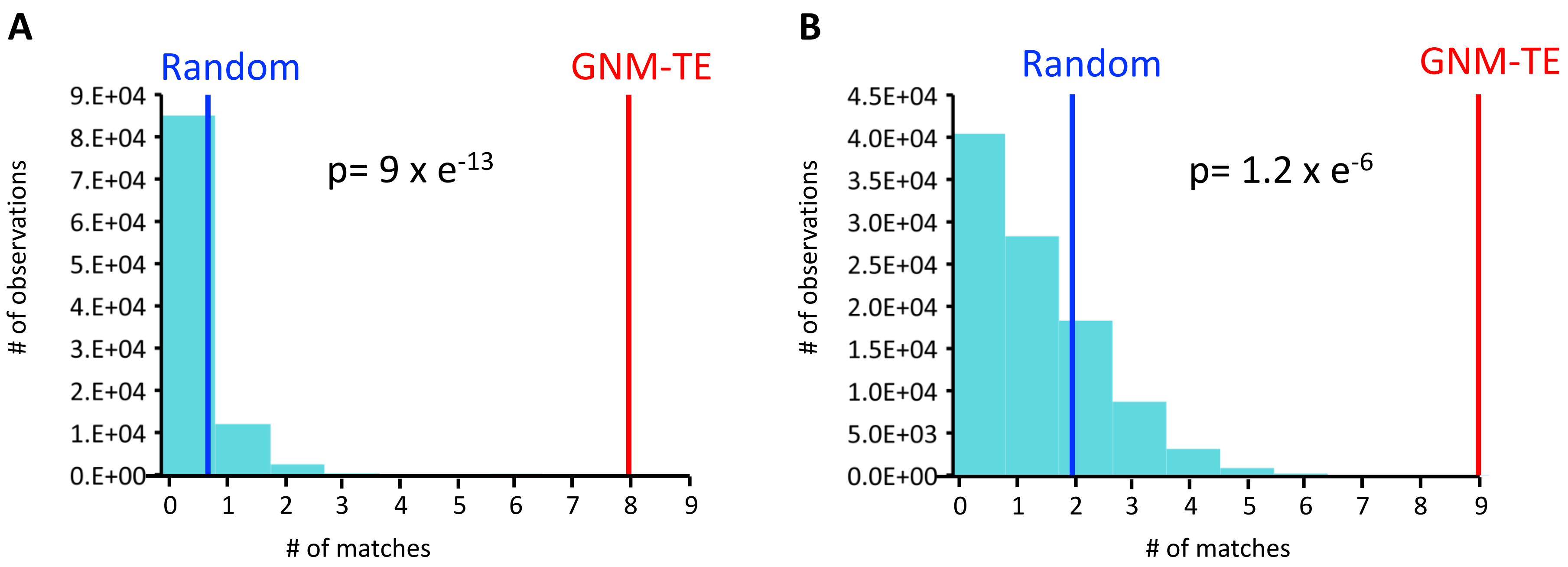


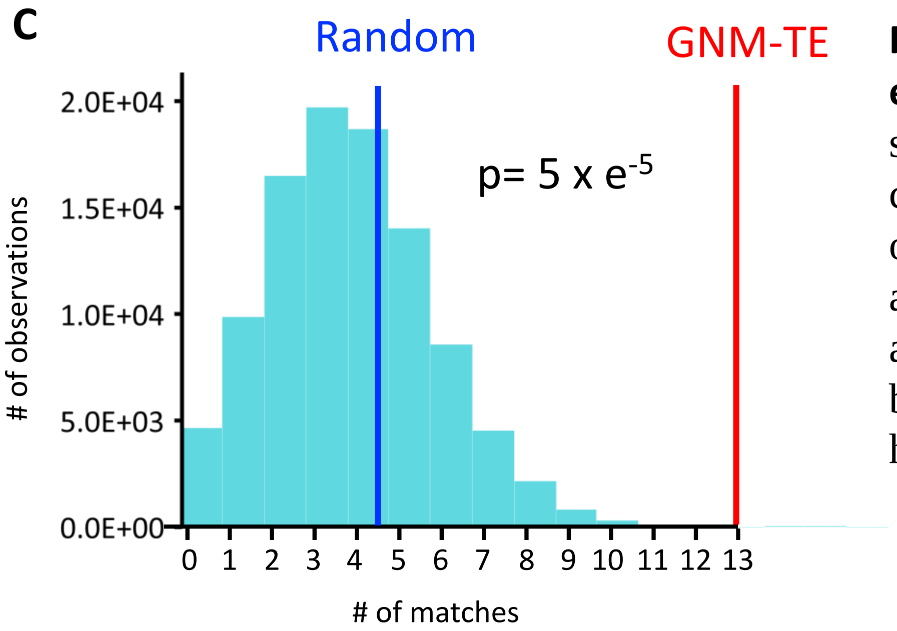


**Figure S4. Correlation between allosteric peaks and positions of the functionally essential residues.** 100,000 sets of 60 randomized positions were generated and for each such set the number of matches with the positions of the 14 essential residues was counted. Shown is the probability distribution when considering only exact matches (A), or also first-coordination sphere interactions within a cutoff distance of ≤ 4 Å (B), or also second-coordination sphere interactions within a cutoff distance of ≤ 7 Å (C). Blue and red vertical lines indicate the mean value of matches for the random and GNM-TE based predictions, respectively. Also shown are the p value obtained by one-tailed hypothesis tests with a significance level of 0.05.

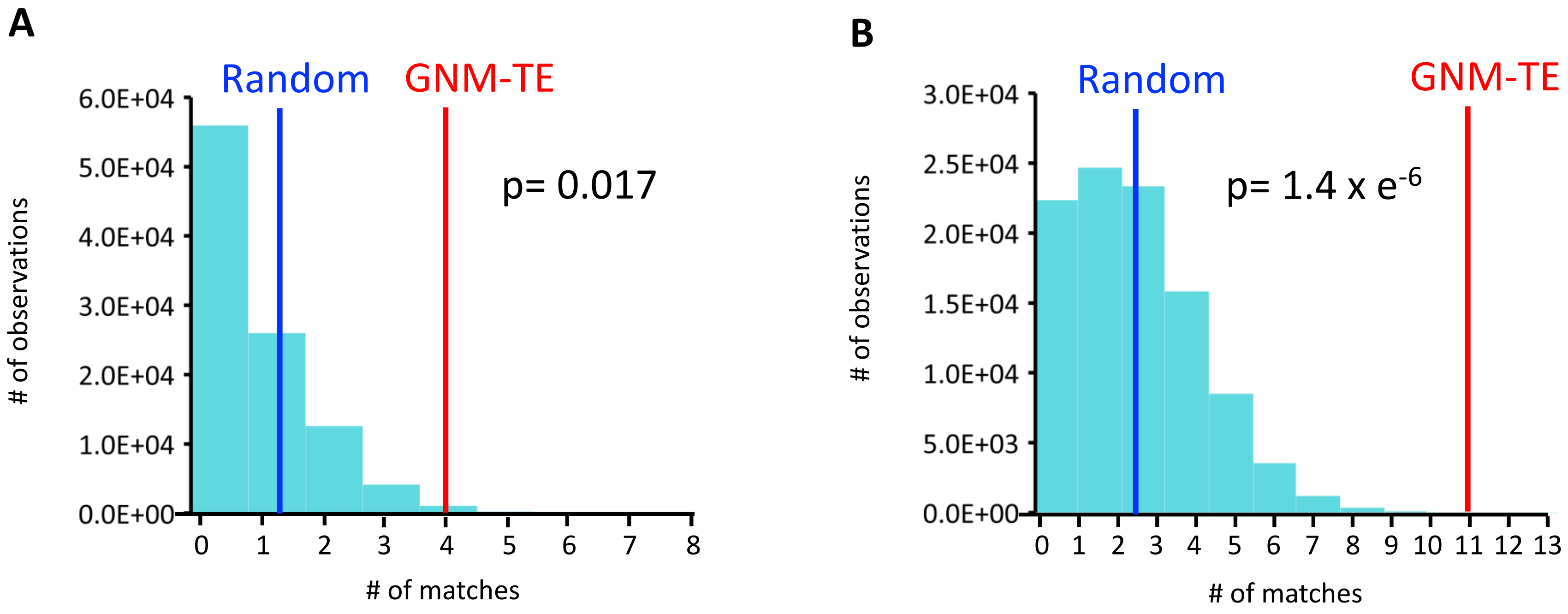


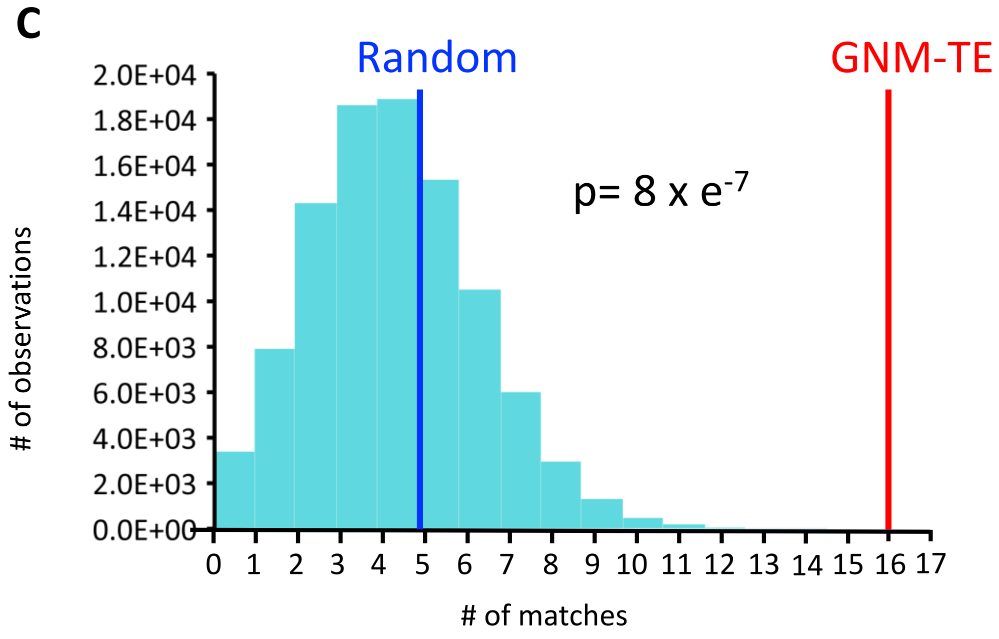


**Figure S5. Correlation between allosteric peaks and ATP binding residues.** 100,000 sets of 60 randomized positions were generated and for each such set the number of matches with the positions of the ATP binding residues was counted. Shown is the probability distribution when considering only exact matches (A), or also first-coordination sphere interactions within a cutoff distance of ≤ 4 Å (B), or also second-coordination sphere interactions within a cutoff distance of ≤ 7 Å (C). Blue and red vertical lines indicate the mean value of matches for the random and GNM-TE based predictions, respectively. Also shown are the p value obtained by one-tailed hypothesis tests with a significance level of 0.05.

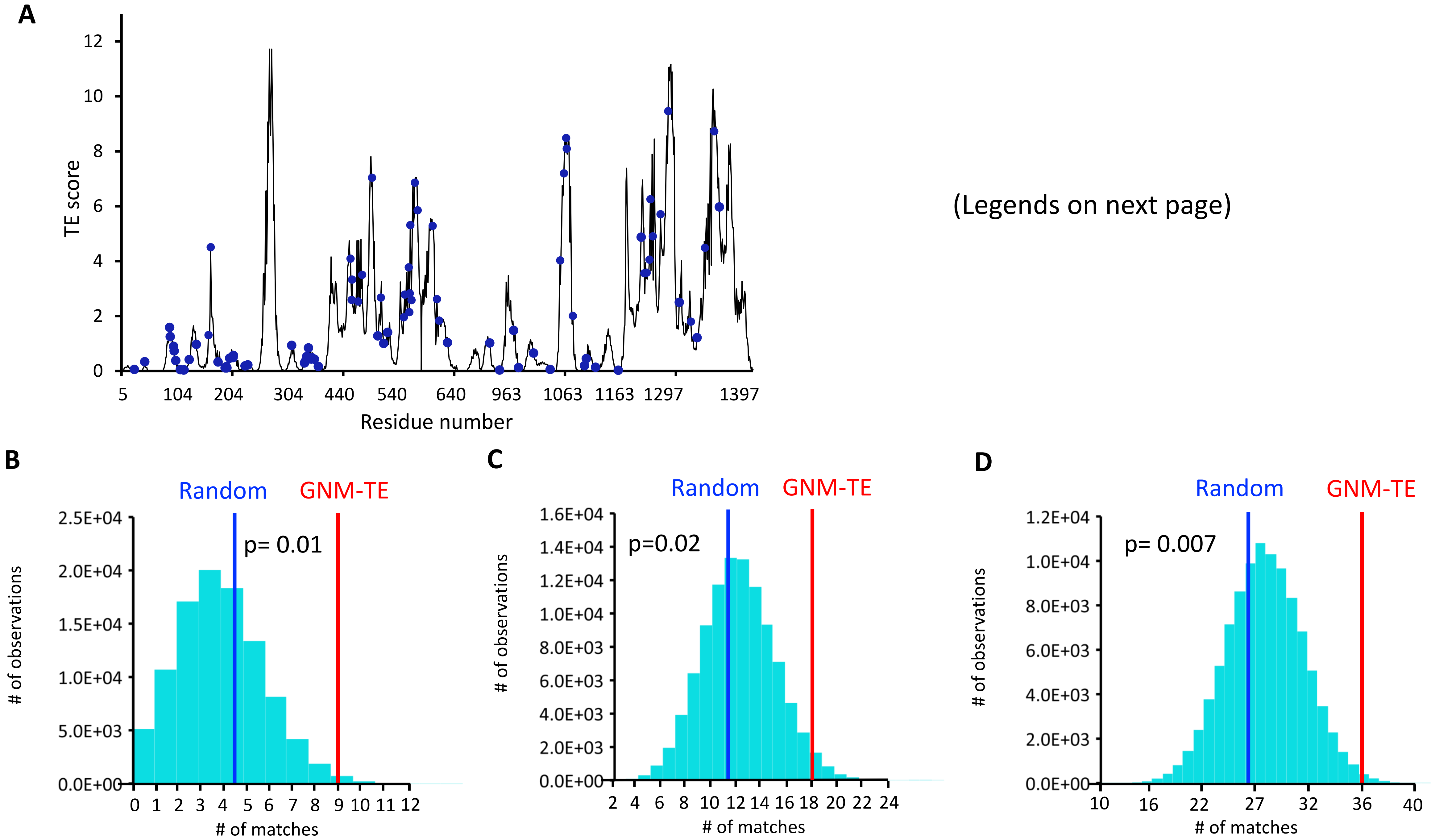


(Legends on next page)

**Figure S6. Correlation between allosteric peaks and positions of disease-causing mutations.**
(A) Shown is the amount of information transmitted (TE score) by each residue of dephosphorylated ATP-free human CFTR (PDB ID 5UAK) calculated using the ten most collective GNM modes (solid black trace). The positions of 86 miss-sense disease causing mutations (https://cftr2.org/) are shown as blue spheres. (B-D) 100,000 sets of 60 randomized positions were generated and for each such set the number of matches with the positions of the disease-causing mutations was counted. Shown is the probability distribution when considering only exact matches (B), or also first-coordination sphere interactions within a cutoff distance of ≤ 4 Å (C), or also second-coordination sphere interactions within a cutoff distance of ≤ 7 Å (D). Blue and red vertical lines indicate the mean value of matches for the random and GNM-TE based predictions, respectively. Also shown are the p values obtained by one-tailed hypothesis tests with a significance level of 0.05.

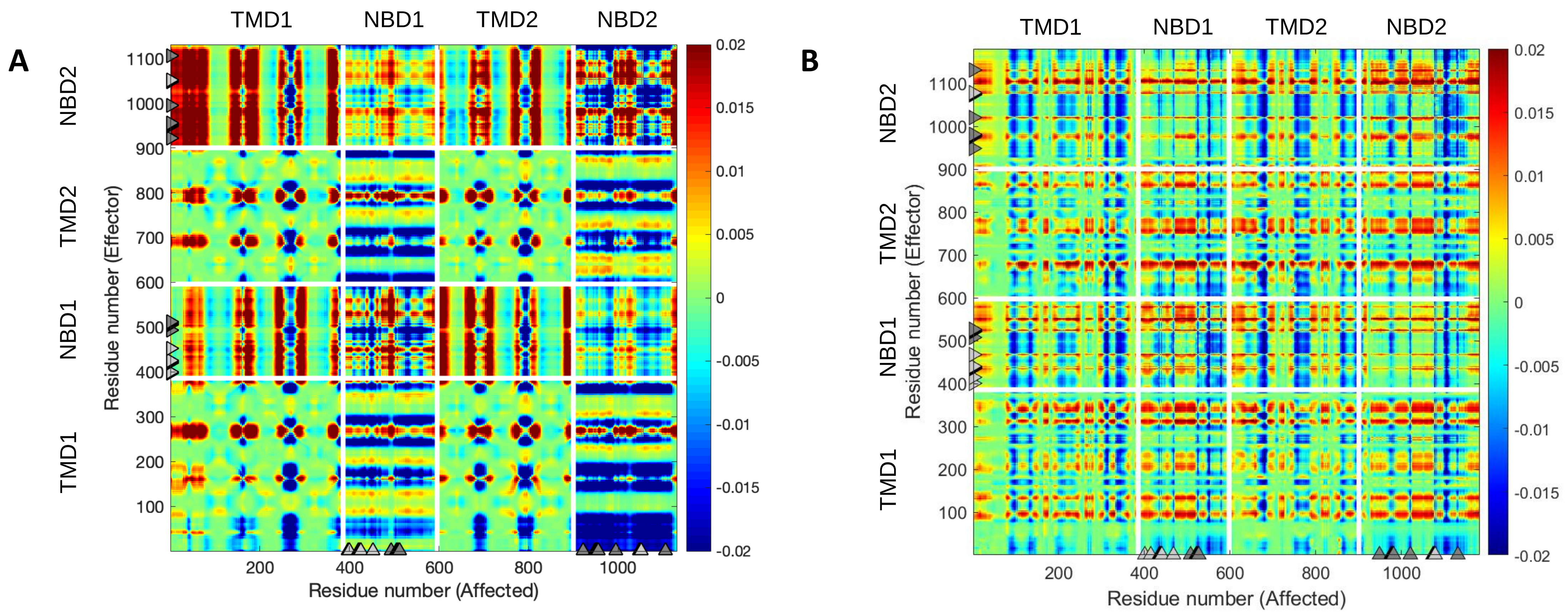


**Figure S7. Phosphorylation and ATP binding rewires the allosteric connectivity in CFTR**

Net transfer entropy (TE) between all residues of CFTR was calculated using the 10 most collective GNM modes for (A) dephosphorylated ATP-free human CFTR (PDB ID 5UAK) and for (B) phosphorylated ATP-bound human CFTR (PDB ID 6MSM). In this 2-D cross correlations map the transfer entropies from effector residues (Y-axis) to the affected residues (X-axis) are color coded, with red and blue colors indicating residues that transmit information (entropy sources) and residues that receive information, respectively. The white lines indicate the domain boundaries, and the grey triangles indicate the locations of the ATP binding sites.

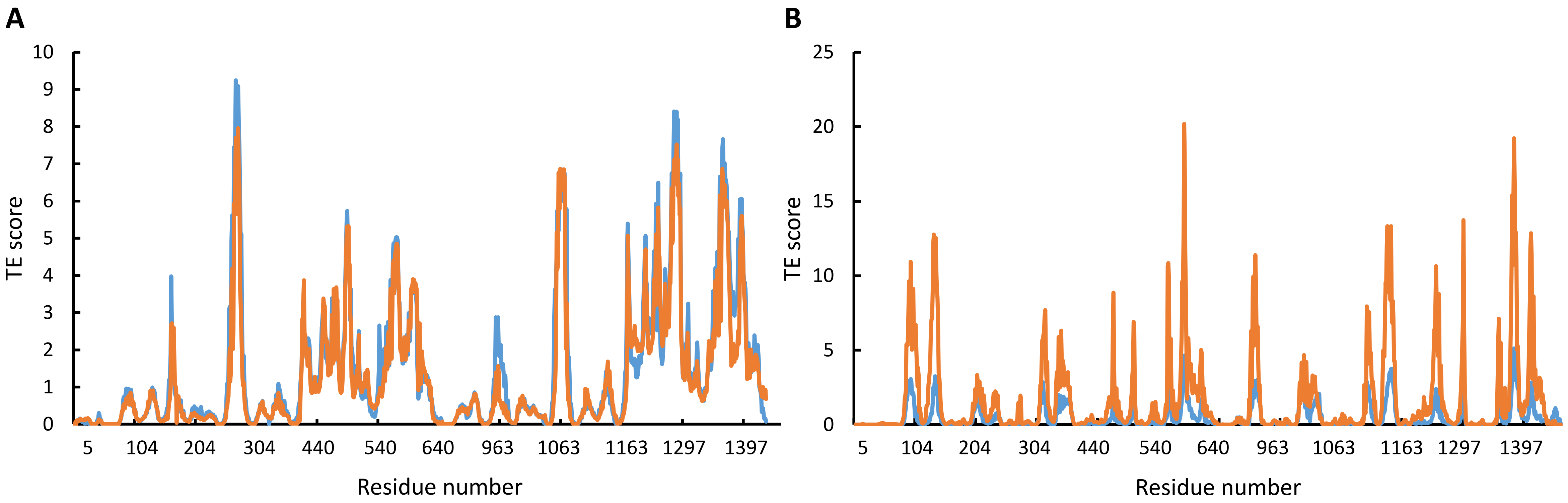


**Figure S8. Conservation of the effect of ATP.** Shown is the amount of information transmitted (TE score) by each residue of dephosphorylated ATP-free (A) or phosphorylated ATP-bound (B) human (blue curves) or zebrafish (orange curves) CFTR (PDB IDs 5UAK, 5UAR, 6MSM, 5W81, respectively).

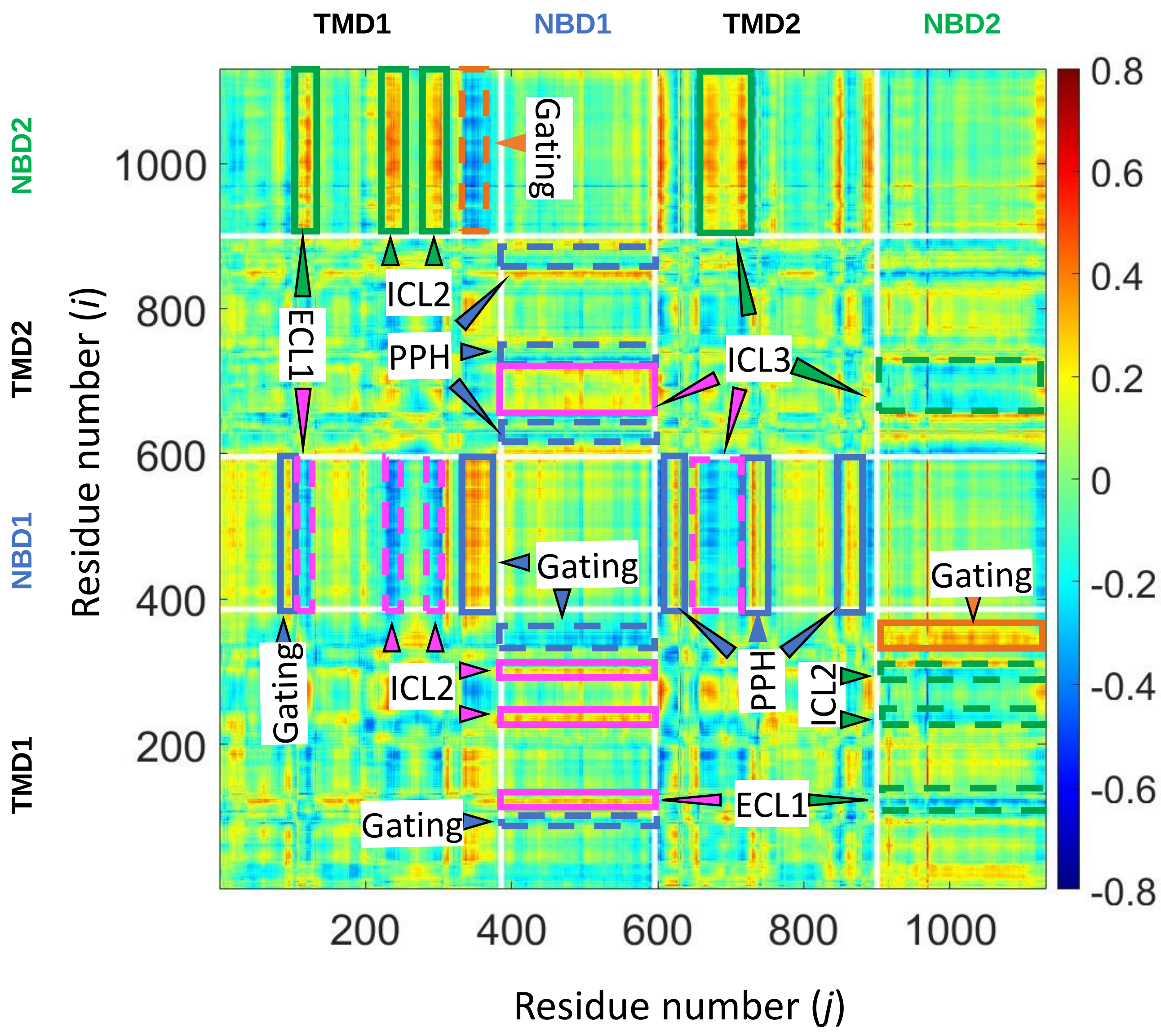


**Figure S9. Directionality of allosteric transduction in CFTR**
Shown is the time-delayed dynamic cross-correlation map for the transition between the ATP-free and ATP-bound conformations of CFTR. A time-delay (τ) of 16 cycles (out of a 50 necessary to complete this transition) was imposed between residues *i* (vertical axis) and residues j (horizontal axis). Red and blue colors represent positive and negative correlations, respectively.
As shown, the synchrony between the movements of NBD2 and ICL2/3 and ECL1 are maintained if NBD2 moves first but are lost if the temporal order of the motions is reversed (compare solid and dashed green rectangles, respectively). Similarly, ECL1 and NBD1 move in synchrony only if the former leads the motion and not *vice-versa* (compare solid and dashed magenta rectangles, respectively), NBD1 leads the motion of the permeation pathway TM helices (PPH, compare solid and dashed blue rectangles, respectively), and the gating residues and the permeation pathway TM helices lead NBD2 (compare solid and dashed orange rectangles, respectively).
